# Norepinephrine inhibits Afferent activation of NTS Catecholamine Neurons in Mice

**DOI:** 10.64898/2026.11.02.763765

**Authors:** Luciano G. Fernandes, Josefina I. Jara, Suzanne M. Appleyard

## Abstract

The nucleus tractus solitarius (NTS) integrates visceral sensory input to regulate feeding behavior, with catecholaminergic (CA) and neuropeptide Y (NPY)-expressing neurons playing key roles in energy homeostasis. Recent evidence suggests functional heterogeneity among CA-NTS neurons, including distinct subpopulations that either suppress or promote feeding. In this study, we investigated how norepinephrine (NE) modulates excitatory synaptic transmission from solitary tract (ST) afferents onto tyrosine hydroxylase-positive (TH⁺) NTS neurons in mice. Using patch-clamp electrophysiology in horizontal brainstem slices, we found that NE inhibits ST-evoked excitatory postsynaptic currents (ST-EPSCs) in a concentration-dependent manner, with maximal effects at ≥10 µM and an EC₅₀ of ∼3.4 µM. This inhibition was mediated by α₂-adrenergic receptors and was independent of GABA_A_ receptor activity. NE also suppressed asynchronous glutamate release and attenuated CCK-induced increase in spontaneous EPSC frequency, suggesting a presynaptic mechanism. NPY similarly reduced ST-EPSC amplitude via Y_2_ receptor activation, and co-application of NE and NPY produce additive or synergistic effects, even at subthreshold concentrations. Together, our results reveal a novel mechanism by which noradrenergic and peptidergic systems interact to regulate visceral sensory processing in the brainstem.

NTS

## Introduction

Vagal afferent fibers transmit afferent visceral signals from the gastrointestinal (GI) system to the central nervous system (CNS), forming their first central synapse in the nucleus tractus solitarius (NTS)(1). These monosynaptic glutamatergic afferents release the excitatory amino acid l-glutamate (2–5) that binds to postsynaptic α-amino-3-hydroxy-5-methyl-4-isoxazolepropionic acid (AMPA) and N-methyl-D-aspartate (NMDA) ionotropic glutamate receptors, thereby activating second-order NTS neurons (5–8) and promoting the integration of the afferent visceral information in the CNS (9–12). As the primary recipient of vagal afferent input, the NTS integrates GI signals and relays this information to both local medullary circuits and higher-order centers involved in energy balance regulation (10, 13, 14), thereby coordinating short-term meal termination and modulating feeding behavior in response to internal physiological states (15–19).

Catecholamine-synthesizing (CA) neurons of the A2 cell group within the NTS (CA-NTS) neurons play a pivotal role in modulating ingestive behavior (20–25). Meal ingestion, gastric distension and satiety signals all activate CA-NTS neurons (26–28), which, in addition to their local actions, promote catecholamine release in several CNS regions, including the lateral parabrachial nucleus (lPBN, Palmiter, 2018; Riche et al., 1990), dorsal motor nucleus of the vagus (DMV, Fukuda et al., 1987; Martinez-Peña y Valenzuela et al., 2004; Sawchenko, 1983; Travagli and Anselmi, 2016), arcuate nucleus of the hypothalamus (Arc, Aklan et al., 2020), and the paraventricular nucleus of the hypothalamus (PVH, Sayar-Atasoy et al., 2023). These projections are well recognized for their roles in regulating feeding behavior (14, 36).

Although various peptides and neurotransmitters modulate vagal afferent neurotransmission onto NTS neurons (Andresen et al., 2012; Appleyard et al., 2007, 2005; Calkins et al., 2025; Cui et al., 2011; Cui et al., 2012a, 2012b, 2012c; Fernandes et al., 2011; Jin et al., 2004b, 2004a; Moran et al., 2001; Winzenried et al., 2025), the effects of norepinephrine (NE) on vagal neurotransmission onto CA-NTS neurons remain poorly understood. Here, we investigated how NE shapes excitatory synaptic transmission onto CA-NTS neurons, with the aim of determining how this modulation influences visceral afferent processing and contributes to the central regulation of feeding behavior.

## Methods

### 1. Animals

Male and female transgenic mice expressing enhanced green fluorescent protein under the control of the tyrosine hydroxylase promoter (TH-EGFP) (48) were maintained on C57BL/6J (N5) background and genotyped for the presence of EGFP by PCR (21). They were housed on a 12 h light/dark cycle at ambient temperature in a murine specific pathogen-free facility of the Vivarium of the Washington State University (Pullman, WA). Chow (Laboratory Rodent Diet no. 5001; PMI Nutrition International, St. Louis, MO) and water were provided *ad libitum*. All animal procedures were conducted with the approval of the Institutional Animal Care and Use Committee in accordance with the U.S. Public Health Service (PHS) Policy on Humane Care and Use of Laboratory Animals (PHS Policy) and the National Institutes of Health (NIH) *Guide for the Care and Use of Laboratory Animals* (NIH Guide).

### 2. Horizontal NTS slices

Hindbrains of TH-EGFP mice (6–16 weeks of age) were prepared as described previously (21, 38). Briefly, slices (250 μm thick) were cut with a sapphire knife (Delaware Diamond Knives, Wilmington, DE) mounted in a vibrating microtome (VT-1000S; Leica Microsystems). Horizontal slices contained a 2–5 mm length of the ST in the same plane as the NTS. Slices were submerged in a perfusion chamber, and all recordings were performed in artificial CSF composed of the following (in mm): 125 NaCl, 3 KCl, 1.2 KH_2_PO_4_, 1.2 MgSO_4_, 25 NaHCO_3_, 10 dextrose, 2 CaCl_2_, bubbled with 95% O_2_/5% CO_2_ at 31.5–32.5°C and pH 7.4 kept at a stable temperature between 31.5°C and 32.5°C. Osmolarity was adjusted to between 305 and 310 mOsm using sucrose. TH+ NTS neurons were identified by EGFP expression and were recorded using patch electrodes filled with a solution composed of the following (in mm): 10 NaCl, 130 K-gluconate, 11 EGTA, 1 CaCl_2_, 2 MgCl_2_, 10 HEPES, 2 NaATP, 0.2 NaGTP, pH 7.3 and 295–299 mOsm. Patch electrodes, 3.8-4.2 MΩ, were guided to neurons using a microscope (Olympus BX51WI) equipped with both fluorescence and differential interference contrast (DIC) optics with infrared (IR) illumination (Figure 1). Voltage-clamp and current-clamp recordings were made with a Multiclamp 700B and pClamp 11 software (Molecular Devices, Union City, CA) with data sampled at 50 kHz and filtered at 3 kHz via an Axon 120 A/D converter and pClamp9 software (Molecular Devices). Only neurons with stable holding currents not exceeding ±100 pA at V_H_ of −60 mV were studied further. Experiments were considered acceptable when a seal resistance of at least 1.0 GΩ was attained, input resistance > 300 MΩ and a series resistance < 20 MΩ that changed less than 10% were observed during recordings. Also, experiments were terminated whenever the holding current reached values higher than +100 pA or lower than -100 pA and this data was not included in the results. No leak subtractions or series resistance compensations were performed. We recorded 190 second-order NTS TH+ neurons (79 from male and 110 from female mice) across 82 animals (43 females, 39 males). Each protocol included at least three animals per sex; in 21 of the 24 experiments, it was more than 3. As no sex-related significant differences were observed in the responses and therefore data from male and female mice were combined for each experiment.

**Figure 1:**
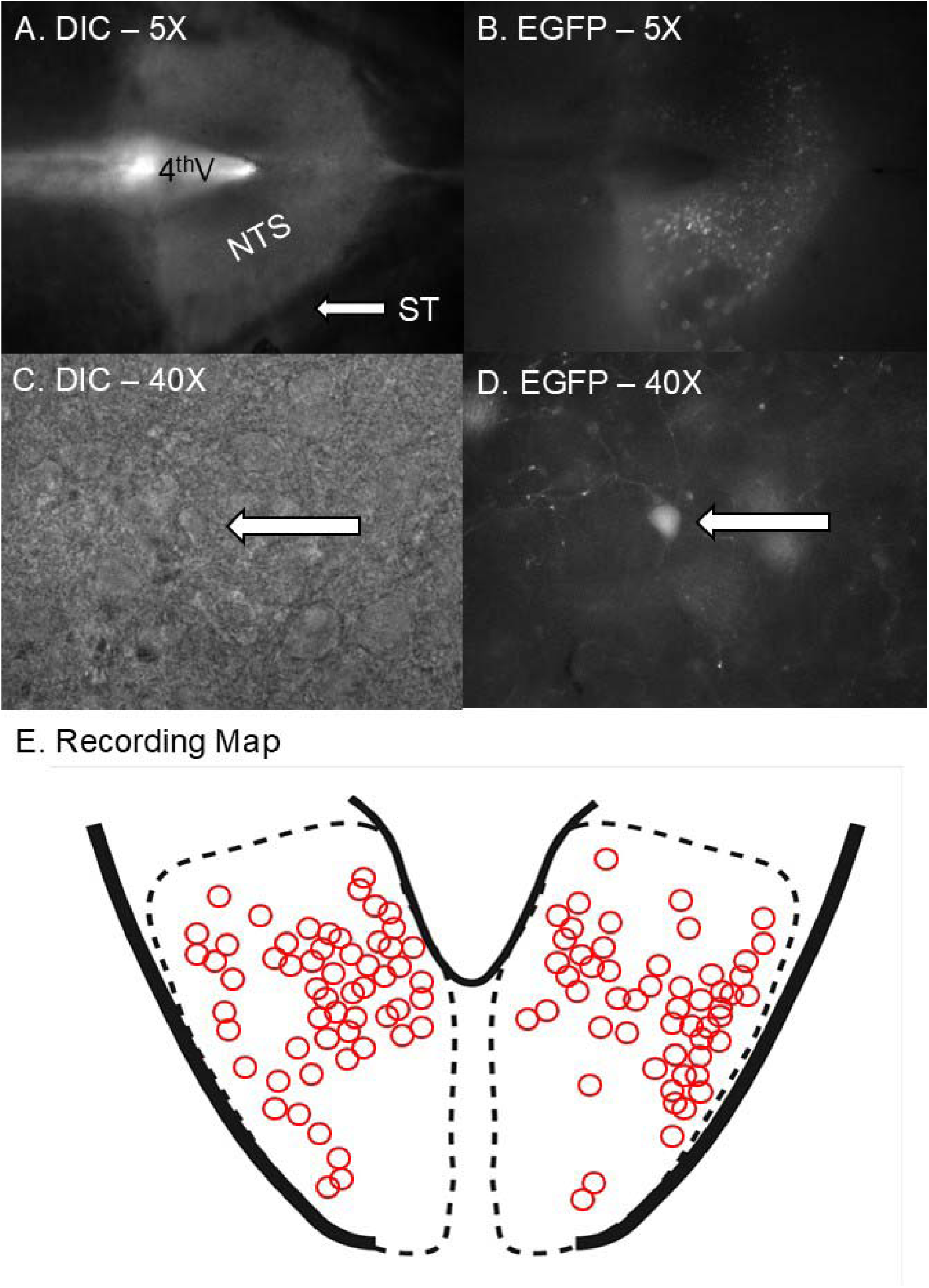
Horizontal brainstem slices contain NTS neurons and the ST at the same anatomical level. Horizontal brainstem slices containing the solitary tract (ST) and the nucleus tractus solitarius (NTS) at the same anatomical level (A) enable direct visualization of NTS neurons using differential interference contrast (DIC) microscopy (B) and identification of tyrosine hydroxylase-expressing neurons in TH-EGFP+ mice (C and D). TH+ neurons were recorded across different NTS levels (E).

### 3. Measurement of synaptic currents

In the present study, we recorded from 165 TH^+^ NTS second-order neurons, from both male and female mice. Evoked excitatory post-synaptic currents (ST-EPSC) were triggered with a concentric bipolar stimulating electrode (F. Haer, Bowdoinham, ME) placed on the ST 1-3 mm from the recording electrode. Bursts of ten shocks (50 Hz) were generated at 6 s intervals and delivered from an isolated programmable stimulator (Master-8; AMPI). Events were collected over 5 minutes in control and then 5 minutes during drug exposure and 5 minutes drug wash out for comparison. Latency was measured for the first of each burst of ten ST-EPSCs in baseline conditions. Jitter was calculated as the standard deviation of 40 ST-EPSCs in baseline conditions and used to classify neurons as either directly contacted by ST afferents (monosynaptic jitter <200 μs, i.e., second order) or by ST polysynaptic pathways (jitter >200 μs, i.e., higher order) (4).

### 4. Spontaneous EPSCs

Spontaneous EPSCs (sEPSCs) were recorded in gap-free mode using voltage clamp. After stabilization of the recordings events were collected over 5 minutes in control and then 5 minutes during drug exposure and 5 minutes drug wash out for comparison. Analysis was performed off-line using pClamp (Molecular Devices) and Mini Analysis (version 5.0; Synaptosoft, Decatur, GA).

### 5. Drugs

The drugs used in the present study were: Norepinephrine (NE 1, 3, 10, 30 µM, Sigma), Phentolamine (PTL, 10 µM, non-selective α adrenergic receptor antagonist, Sigma), SKF86466 (SKF, 15 µM, selective α_2_ adrenergic receptor antagonist, Tocris), Dexmedetomidine (DEX, 10 µM, selective agonist of adrenergic α_2_ receptor, Sigma), Prazosin (PRZ, 10 µM, selective α_1_ adrenergic receptor antagonist, Tocris), Phenylephrine (PLP, 10 µM, selective α_1_ adrenergic receptor agonist, Tocris), Isoproterenol (ISO, 100 µM, non-selective β adrenergic receptor agonist, Tocris), Propranolol (PRO, 100 µM, non-elective β adrenergic receptor antagonist, Tocris), Metoprolol (MET, 100 µM, selective β_1_ adrenergic receptor antagonist, Sigma), BIIE0246 (BIIE, 1 µM, selective Y_2_ receptor antagonist, Tocris) and Neuropeptide Y (NPY, 100 nM, Tocris).

### 6. Statistical analysis

All data are presented as mean ± SEM. Statistical comparisons used either paired or unpaired Student’s *t* test, repeated-measures ANOVA, or one-way ANOVA followed by Tukey posttest and ANOVA two-way. Kolmogorov-Smirnov (K-S) nonparametric analysis was used to compare cumulative distributions of sEPSC amplitudes and frequencies. Values of *p* < 0.05 indicated significant differences. All the statistical analysis, linear correlation, curve fitting graphs and related analysis were conducted by custom made Python (49) scripts, using matplotlib (50) and scipy (51) libraries.

## Results

### 1. Norepinephrine inhibits ST-evoked EPSCs in TH-positive neurons of the NTS in a concentration-dependent manner

We first investigated the concentration-dependent effect of norepinephrine (NE) on the amplitude of the first ST-evoked excitatory postsynaptic current (ST-EPSC_1_) in TH-EGFP^+^ neurons of the nucleus tractus solitarius (NTS). NE application produced a progressive and concentration-dependent reduction in ST-EPSC_1_ amplitude, which conformed to a sigmoidal concentration-response relationship. The mean percentage inhibition at 1 µM, 3 µM, 10 µM, and 30 µM NE was 2.32 ± 2.19%, 2.86 ± 3.05%, 33.77 ± 3.66%, and 31.83 ± 6.12%, respectively (**Figure 2B**). The estimated half-maximal effective concentration (EC₅₀), derived from the fitted curve (R² = 0.993), was 3.43 µM.

**Figure 2.**
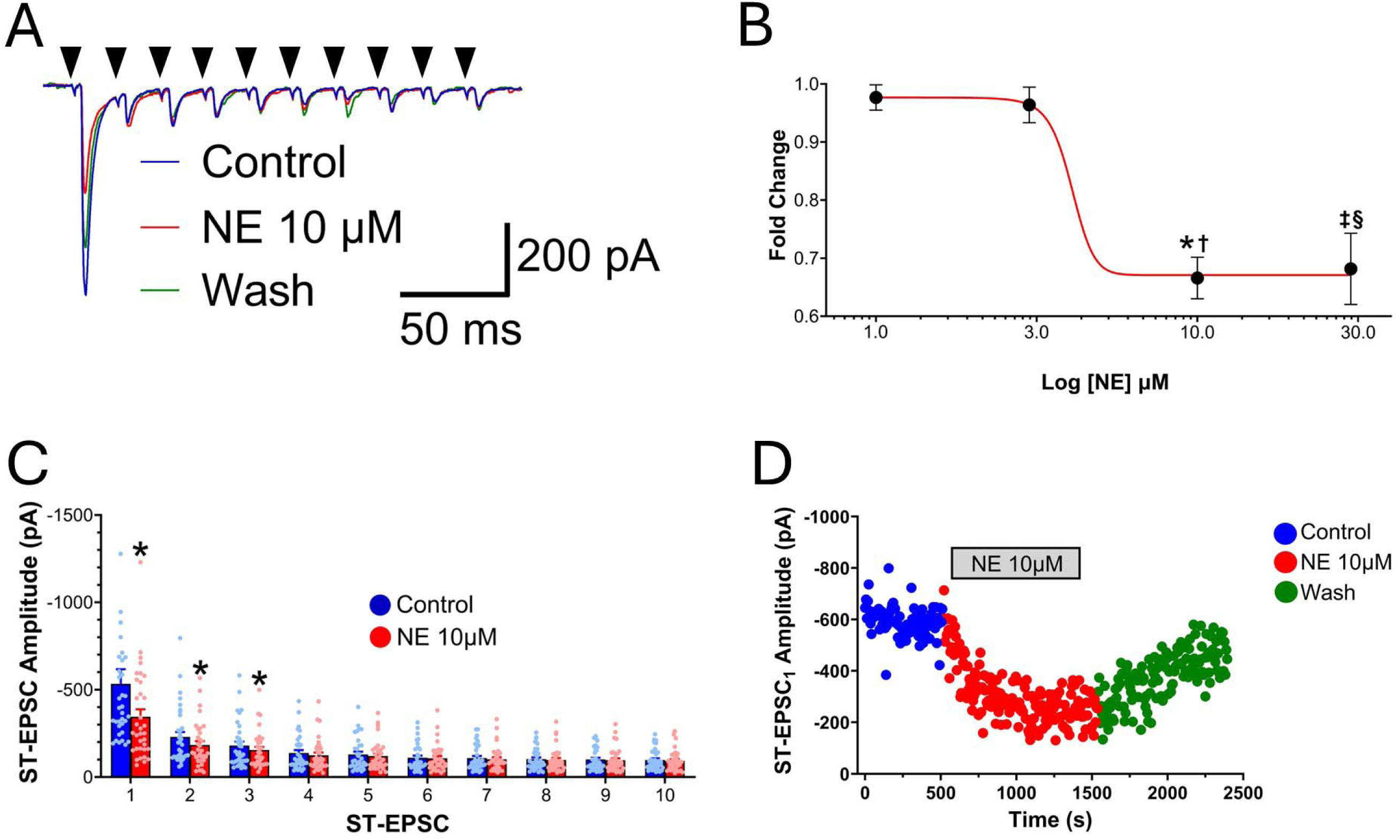
Norepinephrine (NE) inhibits ST-evoked ST-EPSCs in NTS neurons in a dose-dependent manner. Electrical stimulation of the ST (A, black arrowheads) evokes a train of 10 excitatory postsynaptic currents (ST-EPSCs) that exhibit frequency-dependent depression under control conditions (blue trace), during application of NE at 10 µM (red trace), and after washout (green trace). Application of increasing concentrations of NE [B; 1 (n=10), 3 (n=27), 10 (n=34), and 30 (n=16) µM] progressively reduces the amplitude of the first ST-EPSC (ST-EPSC_1_) in a dose-dependent manner (*10 µM vs. 1 µM, p = 0.0001; †10 µM vs. 3 µM, p < 0.0001; ‡30 µM vs. 1 µM, p = 0.0015; §30 µM vs. 3 µM, p < 0.0001). At 10 µM, NE significantly reduces the raw amplitude of the first three ST-EPSCs in the train (C, red bars) compared to control (blue bars; *p < 0.05, same ST-EPSC order). This inhibitory effect of NE on ST-EPSC_1_ amplitude is reversible upon washout (D).

The proportion of NE-responsive neurons (defined as those exhibiting significant inhibition of ST-EPSC_1_ by one-way ANOVA) also increased with concentration. At 1 µM and 3 µM NE, 20.0% (2/10) and 51.9% (14/27) of cells, respectively, were classified as responders. This proportion rose to 88.2% (30/34) at 10 µM and 87.5% (14/16) at 30 µM NE. One-way ANOVA revealed a significant effect of NE concentration on ST-EPSC_1_ inhibition (p = 0.00047, Figure 2B). Post hoc Tukey HSD analysis confirmed that the inhibition at 10 µM NE was significantly greater than at 1 µM (p = 0.0001) and 3 µM (p < 0.00001). Likewise, inhibition at 30 µM NE was significantly greater than at 1 µM (p = 0.0014) and 3 µM (p < 0.00001). No significant difference was observed between 10 µM and 30 µM or between 1 µM and 3 µM concentrations. Collectively, these results demonstrate that NE produces a concentration-dependent inhibition of ST-evoked EPSCs in TH-EGFP+ NTS neurons, with maximal effects observed at concentrations ≥10 µM and an EC₅₀ of approximately 3.43 µM.

When we stimulated the ST 10 times at 50 Hz. we found significant frequency dependent depression, where the amplitude of the evoked EPSC is progressively reduced, as reported previously (Appleyard et al., 2007, **Figure 2A and 2C**). Quantitative analysis confirmed that NE (10 µM) significantly reduced the amplitude of the first three evoked EPSCs. Specifically, ST-EPSC_1_ was reduced from –533.67 ± 84.07 pA (control) to –345.00 ± 43.11 pA during NE application (n = 34, paired t-test, p = 5.95 × 10⁻⁴); ST-EPSC_2_ from –230.01 ± 28.40 pA to –183.97 ± 22.26 pA (p = 1.00 × 10⁻³); and ST-EPSC_3_ from –180.19 ± 22.80 pA to –154.84 ± 17.77 pA (p = 0.021) (**Figure 2C and 2D**). No significant differences were observed for ST-EPSC_4_ through ST-EPSC_10,_ suggesting that the amplitude of the evoked EPSC reaches a steady state.

### 2. Norepinephrine significantly alters Pair Pulse Ratio, but not sEPSC frequency or amplitude in second-order TH-positive NTS neurons

Analysis of the paired-pulse ratio (PPR; ST-EPSC_2_/ST-EPSC_1_) revealed a significant increase during NE application (0.72 ± 0.06) compared to control (0.50 ± 0.03; n = 34, one-way ANOVA, p = 0.0046) (**Figure 3A**), consistent with a presynaptic mechanism of action. However, we did not see any significant difference in either frequency or amplitude when spontaneous excitatory postsynaptic currents (sEPSCs) were recorded from second-order TH-EGFP^+^ neurons of the nucleus tractus solitarius (NTS) under Control, NE 10 µM, and Wash conditions (**Figure 3B**). The mean sEPSC frequency (**Figure 3C**) was 2.9 ± 0.44 Hz (n = 22) in Control, 3.4 ± 0.66 Hz (n = 22) during NE 10 µM application, and 2.7 ± 1.14 Hz (n = 12) during Wash, with a One-way ANOVA for sEPSC frequency revealing no significant effect of treatment (p = 0.8512). Similarly, analysis of sEPSC amplitude showed no significant treatment effect (p = 0.9405, **Figure 3D**). Mean sEPSC amplitudes were 50.03 ± 4.3 pA in Control, 49.09 ± 4.2 pA during NE, and 45.85 ± 5.35 pA in Wash conditions. Cumulative distribution analysis of sEPSC amplitudes (**Figure 3E**) did not reveal a significant difference between Control and NE conditions (KS test, p = 0.502). Likewise, the cumulative distribution of interevent intervals (**Figure 3F**) was not significantly altered by NE application (KS test, p = 0.640).

**Figure 3.**
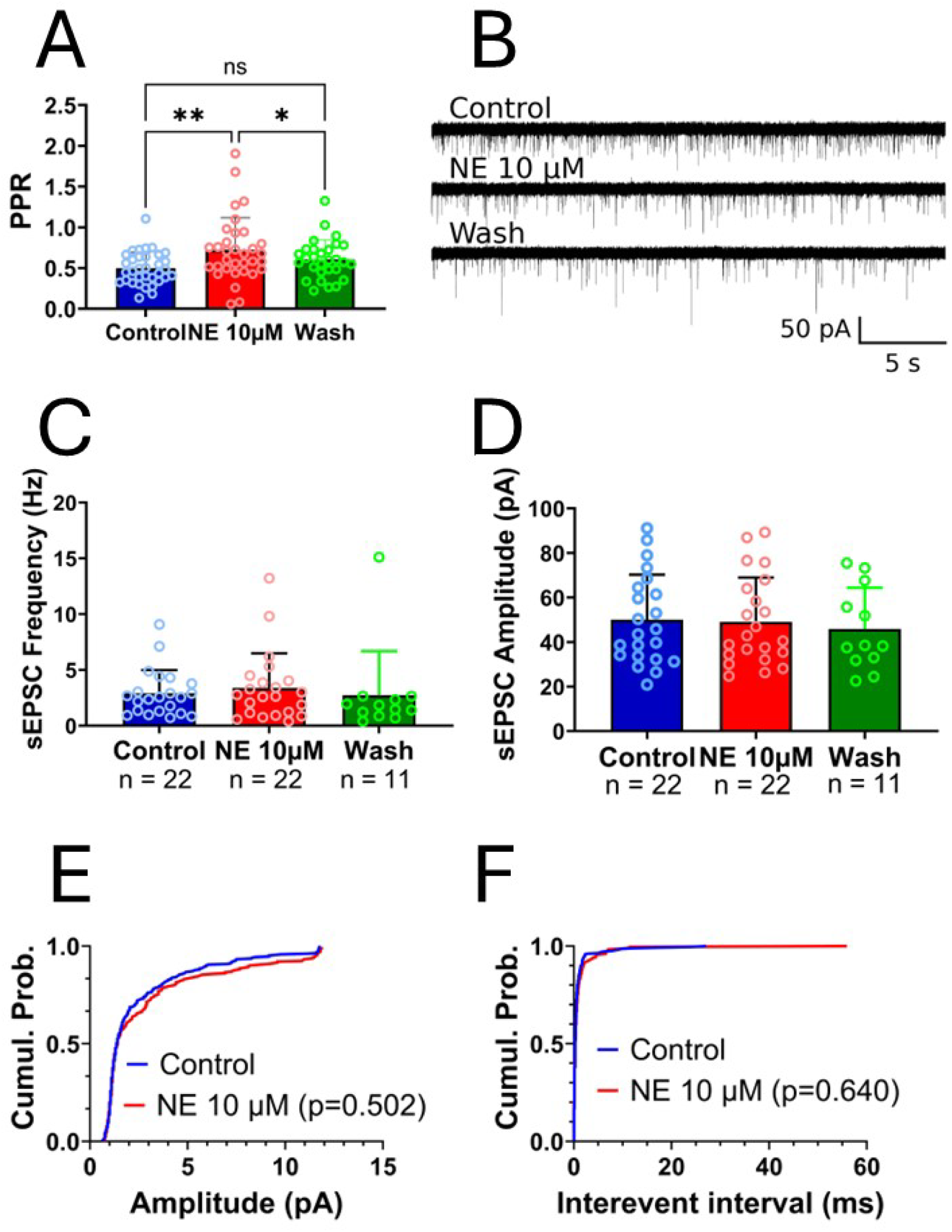
Norepinephrine (NE) reduces the amplitude of ST-EPSCs but does not affect spontaneous EPSC frequency or amplitude. Application of NE (10 µM) significantly reduces the paired-pulse ratio (PPR; ST-EPSC_2_/ST-EPSC_1_) compared to control (A; red bars and scatter vs. blue bars and scatter; **p = 0.0055), consistent with a presynaptic decrease in neurotransmitter release probability. In contrast, NE does not alter the frequency (B, representative traces; C, group data) or amplitude (B, traces; D, group data) of spontaneous excitatory postsynaptic currents (sEPSCs). Cumulative probability plots from a representative cell show no change in sEPSC amplitude (E) or inter-event interval (F) following NE application, further supporting the lack of effect on spontaneous synaptic activity.

### 3. NE reduces asynchronous release from ST terminals

In Figure 4A, spontaneous excitatory postsynaptic currents (sEPSCs) are evident during the initial baseline recording period, followed by evoked EPSCs (ST-EPSCs) elicited by a train of 10 stimuli delivered to the ST at 50 Hz. Immediately following the evoked response, a prominent increase in asynchronous EPSCs (aEPSCs) is observed, which decay rapidly over time, as has been shown previously in other NTS neurons (47, 52–58). This transient increase in aEPSCs following ST stimulation is more clearly visualized in **Figure 4B**, which presents a histogram of aEPSC occurrence from a single cell. The post-stimulation aEPSC frequency shows a marked decline over time when compared to the pre-stimulation baseline. Quantitative analysis of the aEPSC frequency is shown in **Figure 4C**. NE significantly decreased the raw peak aEPSC frequency (82.96 ± 15.09 Hz, n = 27) compared to control conditions (168.8 ± 34.04 Hz, n = 27; p = 0.0031, t-test). When normalized to the baseline period, NE also significantly reduced the aEPSC frequency (3.51 ± 0.47 Hz, n = 27) relative to control (5.04 ± 0.47 Hz, n = 27; p = 0.0211, t-test), as shown in **Figure 4D**. In contrast, NE had no significant effect on the mean amplitude of aEPSCs (**Figure 4E**, NE: 49.39 ± 3.94 pA vs. Control: 48.46 ± 4.01 pA; p = 0.5495, t-test).

**Figure 4.**
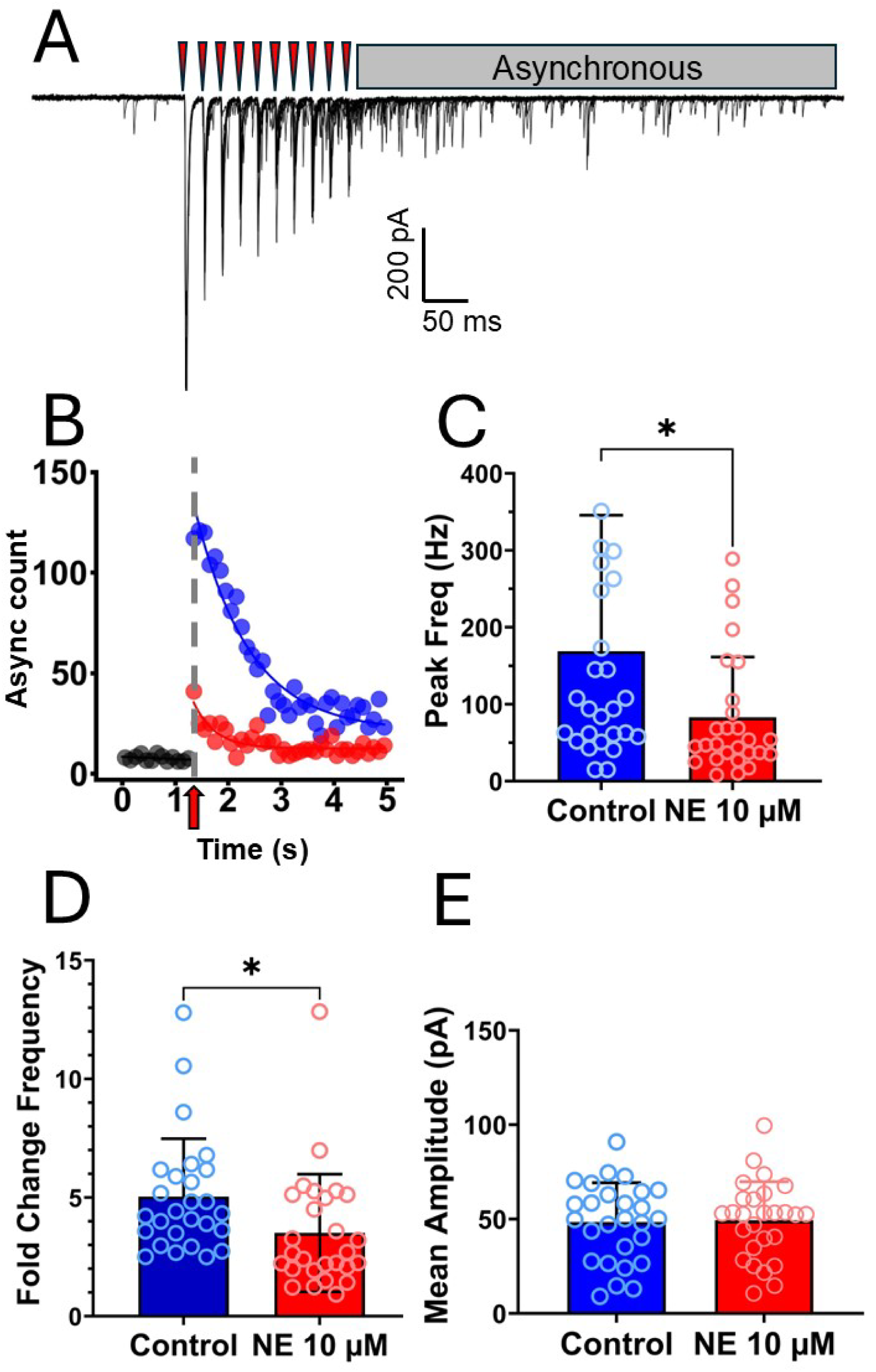
Norepinephrine suppresses asynchronous glutamate release from solitary tract terminals onto TH^⁺^ NTS neurons. **(A)** Representative trace from a TH^⁺^ NTS neuron showing superimposed responses from 10 sweeps. A train of 10 stimuli delivered at 50 Hz (red arrowheads) evokes synchronous EPSCs (ST-EPSCs). Following the ST-EPSCs, a prolonged period of asynchronous glutamate release is evident. **(B)** Histogram (100 ms bins) of aEPSC events from a single neuron, illustrating baseline activity (first second) and the post-ST stimulation period (red arrow), where asynchronous release decays over time. Data are shown before (Control, blue scatter) and after NE 10 µM application (NE, red scatter). **(C)** Group data showing the peak aEPSC frequency following ST stimulation before (Control, blue bar) and after NE 10µM treatment (NE, red bar; *n* = 22 per group). **(D)** Mean aEPSC frequency normalized to baseline, demonstrating a significant reduction following NE application (Control, blue bar; NE, red bar; *n* = 22 per group). **(E)** Mean aEPSC amplitude before (Control, blue bar) and after NE 10µM superfusion (NE, red bar; *n* = 22 per group), showing no significant change.

### 4. α₂-Adrenergic Receptors Mediate the Inhibitory Effect of NE on ST-EPSC amplitude

To delineate the receptor mechanisms underlying NE-mediated suppression of excitatory transmission, we employed selective agonists and antagonists targeting adrenergic receptor subtypes. Fold change analysis revealed that bath application of the selective α₂-adrenergic receptor agonist dexmedetomidine (DEX, 10 µM) closely replicated the inhibitory effect of NE (10 µM) on ST-EPSC_1_ amplitude (NE: 0.67 ± 0.03, *n* = 35; DEX: 0.70 ± 0.04, *n* = 9; one-way ANOVA, *p* = 0.9874; **Figures 5A and 5C**).

**Figure 5.**
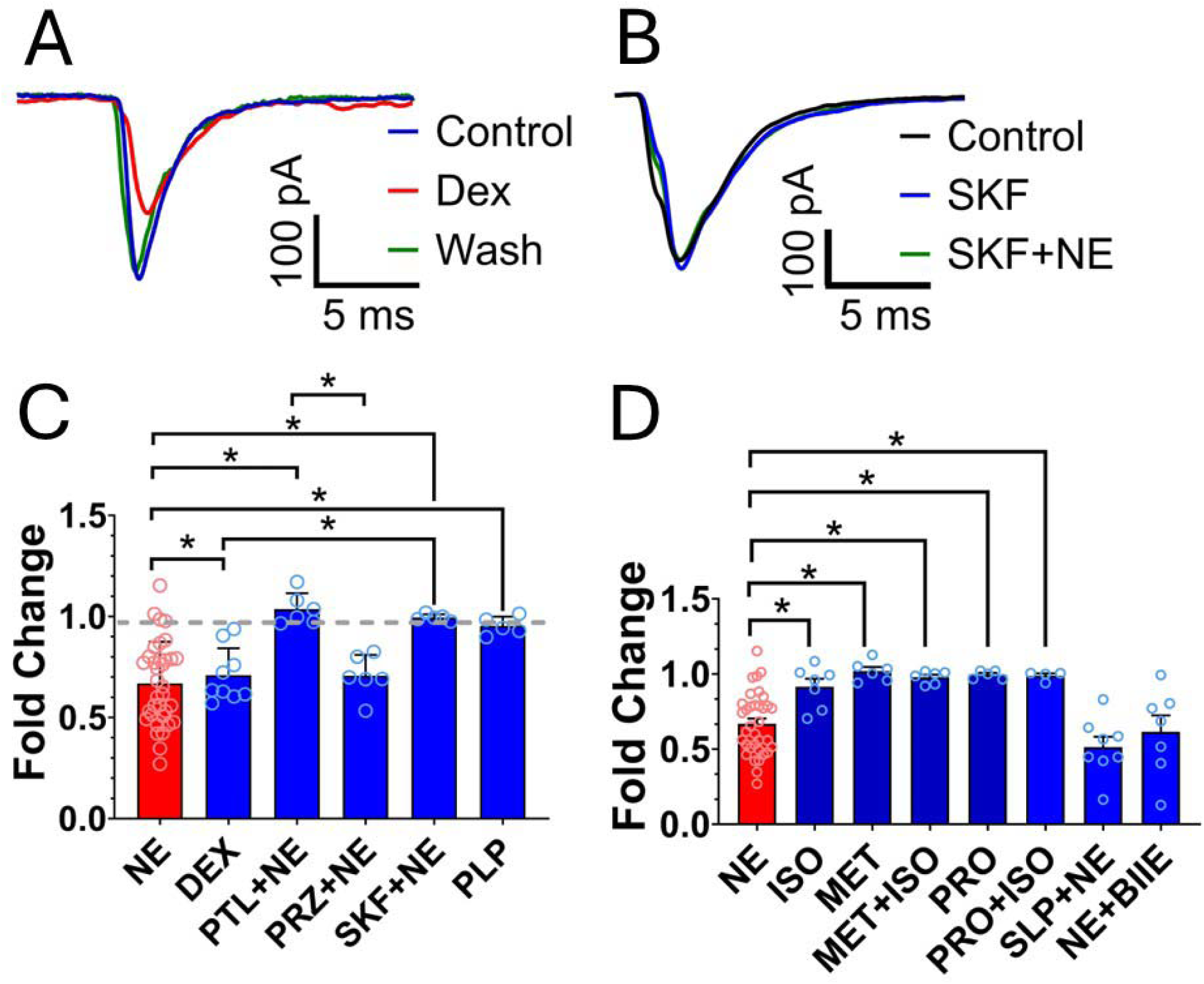
α_2_-adrenergic receptors mediate the inhibitory effect of norepinephrine (NE) on ST-EPSC_1_ amplitude. Application of the selective α_2_-adrenergic receptor agonist dexmedetomidine (10 µM, n=9) mimics the inhibitory effect of NE (10 µM, n=35) on the amplitude of the first evoked EPSC (fold change of ST-EPSC_1_), as shown in the representative trace (A, red) and group data (C, DEX). Co-application of NE with the non-selective α-adrenergic antagonist phentolamine (10 µM; B, PTL+NE, n=6) blocks the effect of NE, indicating a role for α-adrenergic receptors. This blockade is not observed with the selective α_1_ antagonist prazosin (10 µM; C, PRZ+NE, n=6) but is replicated by the selective α_2_antagonist SKF86466 (15 µM; B and C, SKF+NE, n=5), supporting the involvement of α_2_ receptors. Activation of α_1_ receptors using the selective agonist phenylephrine (10 µM; C, PLP, n=5) does not mimic the effect of NE. Furthermore, the application of the non-selective β-adrenergic agonist isoproterenol (100 µM; D, ISO, n=7), the non-selective β antagonist propranolol (100 µM; D, PRO, n=5), and the selective β_1_ antagonist metoprolol (100 µM; D, MET, n=6) fail to alter ST-EPSC_1_ amplitude. The D_2_ receptor antagonist (20 µM; D, SLP, n=8) and the Y_2_ receptor antagonist (1 µM, D, BIIE, n=7) failed to block the inhibitory effect onto ST-EPSC_1_ amplitude. *p < 0.05, **p < 0.01, ***p < 0.001.

Furthermore, the effect of NE on ST-EPSC amplitude was prevented by co-application of NE with the non-selective α-adrenergic receptor antagonist phentolamine (10 µM) (PTL+NE: 1.03 ± 0.03, *n* = 6; *p* = 0.0001), implicating an α-adrenergic receptors in mediating this response (**Figure 5C**). Co-application of the α₂-adrenergic receptor antagonist SKF86466 and NE prevented the depression on ST-EPSC amplitude promoted by NE (NE: 0.67 ± 0.03, *n* = 35; SKF: 0.99 ± 0.007, *n* = 5; one-way ANOVA, *p* = 0.0018; **Figures 5B and 5C**), but the blockade of α₁ receptors with the specific α_1_ receptor antagonist prazosin (10 µM) did not prevent the inhibitory effect of NE (PRZ+NE: 0.70 ± 0.04, *n* = 6; *p* = 0.9956), confirming the specific involvement of α₂ receptor activation for NE’s action. Moreover, activation of α₁ receptors with the selective agonist phenylephrine (10 µM; PLP: 0.97 ± 0.02, *n* = 5) failed to replicate NE’s effect (*p* = 0.0092; **Figure 5C**), further underscoring the specificity of the α₂ receptor pathway.

To assess the involvement of β-adrenergic receptors, we tested multiple pharmacological agents targeting this class. The non-selective β agonist isoproterenol (100 µM; ISO: 0.91 ± 0.05, *n* = 7; *p* = 0.0064), the non-selective β antagonist propranolol (100 µM; PRO: 0.99 ± 0.012, *n* = 5; *p* = 0.0019), and the selective β₁ antagonist metoprolol (100 µM; MET: 1.02 ± 0.03, *n* = 6; *p* = 0.0002) all failed to alter ST-EPSC_1_ amplitude in comparison to NE alone (0.67±0.03, n=35, **Figure 5D**). Furthermore, combined application of propranolol and isoproterenol (PRO+ISO: 0.99 ± 0.015, *n* = 4; *p* = 0.0081) or metoprolol and isoproterenol (MET+ISO: 0.99 ± 0.017, *n* = 6; *p* = 0.0013) did not block the effects of NE on the ST-EPSC amplitude (**Figure 5D**).

The co-application of the specific Dopaminergic D_2_ receptor antagonist sulpiride and NE (SLP+NE, 0.51±0.07, **Figure 5D**, n=8, p=0.3894) failed to block the inhibitory effect of NE on the ST-EPSC_1_ amplitude in TH+ NTS neurons, indicating that the effect is not mediated by D_2_ receptors. Lastly, the co-aplication of the specific Y_2_ receptor antagonist BIIE 0246 and NE also does not affect the inihibitory effect of NE onto ST-EPSC_1_ (NE+BIIE, 0.61±0.11, **Figure 5D**, n=7, p = 0.9966).

Collectively, these data demonstrate that the inhibitory effect of NE on ST-evoked EPSCs is mediated specifically by α₂-adrenergic receptors and does not involve α₁-, β-adrenergic, Y_2_ or D_2_ receptor activation.

### 5. NE and NPY promote ST-EPSC depression in an additive manner

To investigate potential interactions between norepinephrine (NE) and neuropeptide Y (NPY) in modulating excitatory neurotransmission in the nucleus tractus solitarius (NTS), we performed voltage-clamp recordings of ST-EPSCs in second-order NTS neurons. Analysis of ST-EPSC_1_ **fold change** (**Figure 6A**) revealed no significant difference among the effects of NE (10 µM; 0.61 ± 0.03, *n* = 20), NPY (100 nM; 0.66 ± 0.05, p=0.9126, *n* = 18), but both are significantly different from their co-application (0.45 ± 0.042, *n* = 21, p=0.0158 from NE and p=0.0020 from NPY). Although, **paired-pulse ratio (PPR)** analysis (**Figure 6B**) showed no significant difference among NE (0.79 ± 0.07, p=0.76, n=20) and NPY (0.94 ± 0.38, p= 0.23, n=18), and NE+NPY co-application (1.23 ± 0.28, n=21) was different from both NE (p = 0.0158) and NPY (p = 0.002) applications. In the PPR analysis, all groups tested were different from control (p=0.0004, n=21). Taken together this data suggests that NE and NPY mechanisms do interact to present additive effect further depressing ST-EPSC amplitude. The co-application of NE and NPY antagonists (SKF and BIIE, respectively) did not affect ST-EPSC_1_ fold change (0.89 ± 0.21, n = 21, Figure 6A) or PPR (0.69 ± 0.47, Figure 6B) in relation to controls.

**Figure 6.**
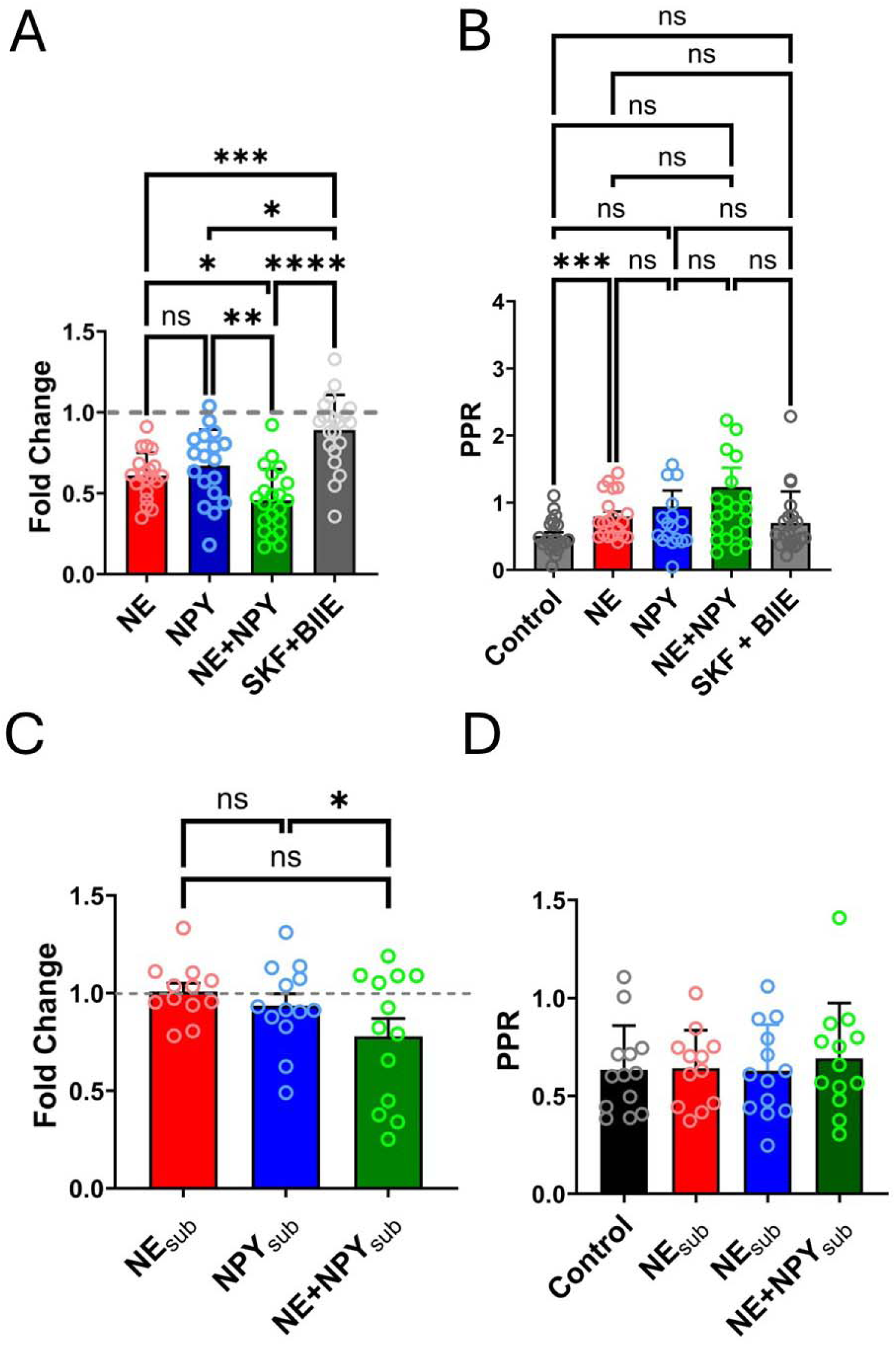
Norepinephrine (NE) and neuropeptide Y (NPY) exert additive effects on ST-EPSC_1_ inhibition. Co-application (n=18) of NE (10 µM) and NPY (100 nM) produced a greater reduction in ST-EPSC_1_ amplitude (A, Fold Change) but not in the paired-pulse ratio (B, PPR = ST-EPSC_2_/ST-EPSC_1_) compared to either compound alone (NE n=20, NPY n=17). Similarly, co-application of lower concentrations (n=13) of NE (3 µM) and NPY (30 nM) also produced an enhanced inhibitory effect on ST-EPSC_1_ amplitude (C, Fold Change) but not in paired-pulse ratio (D, PPR) in comparison to each compound isolated (NE n=12, NPY n=12). Co-application of α_2_ and Y_2_ antagonists did not change ST-ESPC1 amplitude (A) or PPR (B). *p < 0.05, **p < 0.01, ***p < 0.001.

We next determined whether subthreshold concentrations of NE and NPY might interact synergistically, we applied lower concentrations of each compound [NE_sub_ (3 µM) and NPY_sub_ (30 nM)] alone or in combination. **Fold change** analysis of ST-EPSC_1_ amplitude (**Figure 6C**) showed that the co-application of NE_sub_ and NPY_sub_ (n=13) produced a summation effect in respect to NPY_sub_ alone (n=13, p=0.0362). No significant difference between NE 3 µM (1.00 ± 0.042, *n* = 12) and NPY 30 nM (0.93 ± 0.06, p=0.43, *n* = 13), and in comparison to control were found, suggesting that these subthreshold concentrations do interact to produce a greater inhibitory effect at presynaptic level, even though isolated application of subthreshold concentrations do not change ST-EPSC amplitude. On the other hand, **PPR** analysis (**Figure 6D**) revealed no significant differences across the same conditions: NE 3 µM (0.64 ± 0.05, p=0.98, n=12), NPY 30 nM (0.62 ± 0.06, p=0.99, n=13), and NE+NPY (0.72 ± 0.07, p=0.5, n=13) in comparison to control (0.63±0.062, n=13).

To assess whether the effects of NE or NPY on ST-NTS transmission depend on GABA_A_receptor activation, we examined ST-EPSC_1_ amplitude and PPR in the presence or absence of the GABA_A_ receptor antagonist gabazine (5 µM). Fold change analysis (**Figure 7A**) showed that both NE (10 µM; 0.60 ± 0.035, *n* = 10) and NPY (100 nM; 0.69 ± 0.08, *n* = 11) significantly reduced ST-EPSC_1_ amplitude compared to gabazine alone (0.96 ± 0.03, *n* = 20; one-way ANOVA, NE vs. gabazine: *p* = 0.0015; NPY vs. gabazine: *p* = 0.016), but did not differ from each other (*p* = 0.8968). Co-application of gabazine and NE also reduced ST-EPSC_1_ amplitude (0.71 ± 0.083, *n* = 10), an effect significantly different from gabazine alone (*p* = 0.0453) but not from NE alone (*p* = 0.8485).

**Figure 7.**
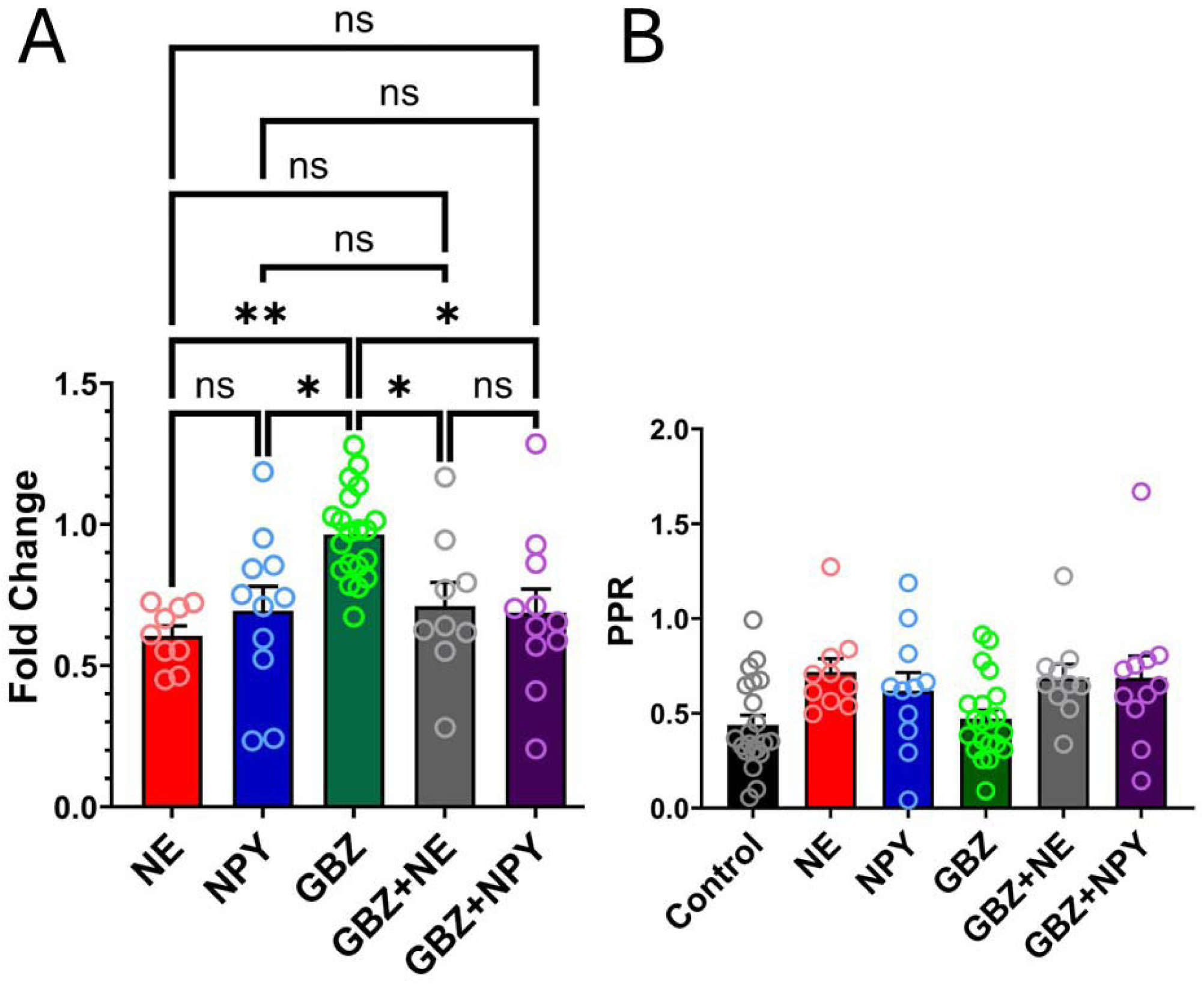
The inhibitory actions of NE (10 µM) and NPY (100 nM) are not mediated by GABA_A_ receptor activation at ST terminals. The co-application of NE 10 µM (n=9) or NPY 100 nM (n=11) with the GABA_A_ receptor antagonist gabazine 5 µM does not alter the depressive effects of NE (A, GBZ+NE, n=9) or NPY (A, GBZ+NPY, n=11). Both groups were similar to NE and NPY alone and different from the isolated application of gabazine (n=20). No effect was observed in PPR following the application of NE, NPY, gabazine or combined application of gabazine with NE or NPY (B). *p < 0.05; **p < 0.01.

**PPR** analysis (**Figure 7B**) showed that NE (10 µM) did not increase PPR (0.72 ± 0.071, n = 10) compared to control (0.43 ± 0.067, n = 21; p = 0.0756). Co-application of gabazine and NE (0.69 ± 0.072, n = 10) similarly did not alter PPR relative to control (p = 0.1489), and values were not different from NE alone (p = 0.9998). A similar pattern was observed with gabazine during NPY application: the effect of NPY alone (0.62 ± 0.09, n = 11) did not differ from the co-application of NPY + gabazine (0.69 ± 0.11, n = 11), from gabazine alone (0.47 ± 0.04, n = 21), or from control (0.43 ± 0.06, n = 21).

Thus, these findings demonstrate that NE and NPY reduce excitatory synaptic transmission at ST-NTS synapses in an additive manner and that these effects are independent of GABA_A_ receptor-mediated inhibition. Furthermore, because simultaneous blockade of both α_2_ and Y_2_ receptors produced no detectable effect on ST-EPSC_1_ amplitude or PPR, these findings suggest the absence of a tonic influence of norepinephrine (NE) and neuropeptide Y (NPY) on ST terminals.

### 6. NE reduced CCK-induced increase in spontaneous glutamate release

We have previously shown that NPY can inhibit the effects of CCK on sEPSC frequency (39). We therefore next evaluated whether a preapplication of NE inhibits the effects of CCK (100nM CCK) on the amplitude and frequency of spontaneous events. pre- and co-application of NE reduced the frequency of sEPSC release induced by CCK both in the peak (fold change: CCK=3.18±0.6 vs CCK+NE=1.33±0.32, p=0.00661, n=9, **Figure 8B**) and in the steady-state phase (fold change: CCK=2.5±0.35 vs CCK+NE=1.89±0.35, p=0.024, n=9, **Figure 8C**). This result indicates that NE also affects sEPSC release induced by CCK.

**Figure 8.**
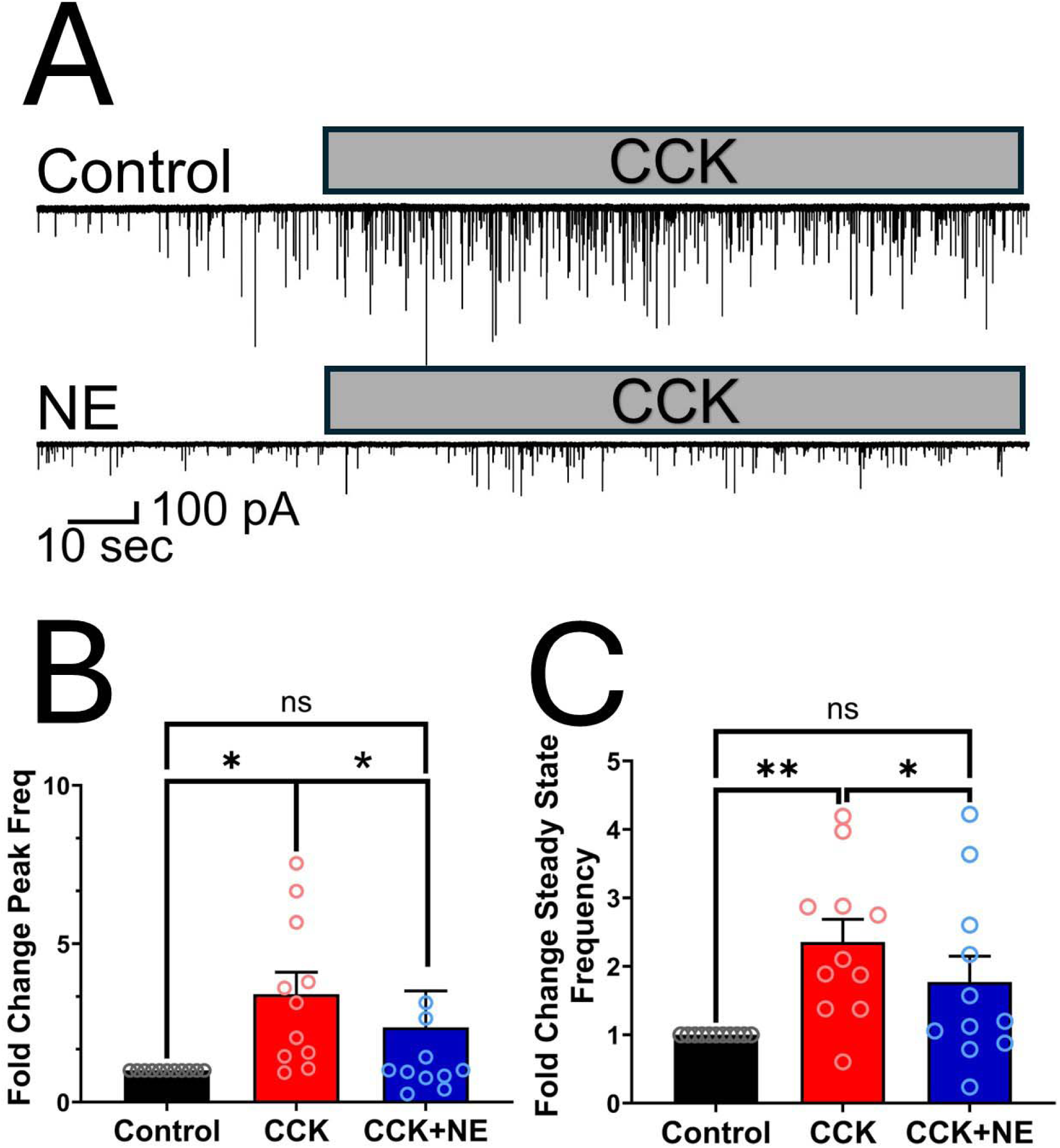
Norepinephrine (NE) attenuates CCK-induced enhancement of sEPSC frequency. Superfusion with cholecystokinin (CCK, 100 nM) significantly increases the frequency of spontaneous excitatory postsynaptic currents (sEPSCs, panel A top trace and panels B and C red bars, n=8) in comparison to control (panels B and C, black bars, n=8). Co-application of NE 10 µM and CCK 100 nM (A, bottom trace) reduces the normalized peak frequency of sEPSC (B, blue bars, n=8) and normalized steady-state frequency (c, blue bars), in comparison to control and to CCK alone. *p < 0.05; **p < 0.01.

## Discussion

Norepinephrine inhibits glutamatergic transmission evoked by ST stimulation onto identified TH^+^ NTS neurons through an α_2_ adrenergic mechanism. This inhibition was accompanied by an increased paired pulse ratio and reduced asynchronous glutamate release, supporting a presynaptic component of NE action. Although NE did not measurably alter basal sEPSC frequency or amplitude, it attenuated the increase in sEPSC frequency elicited by CCK. Combined application with NPY further strengthened the inhibition of transmission evoked by ST stimulation. Together, these findings identify a novel mechanism through which noradrenergic and peptidergic signalings interact to reduce visceral afferent drive to NTS catecholaminergic neurons and possibly modify their responses to signals related to feeding.

NE inhibition of ST-EPSC amplitude increased with concentration and was observed in most neurons at 10 to 30 µM. The effect therefore appears broadly distributed within the TH^+^ population sampled here. The pharmacological results support the involvement of α_2_ receptors, as dexmedetomidine reproduced the inhibition and both nonselective α adrenergic blockade and selective α_2_ antagonism prevented it. Moreover, α_1_ blockade did not prevent NE-induced inhibition, and α_1_ receptor activation did not reproduce it. Neither D_2_ nor Y_2_ receptor blockade prevented the NE response. Thus, we identified α_2_ receptors as the principal mediators of inhibition of evoked EPSCs by NE under our recording conditions.

This receptor profile extends earlier works on adrenergic regulation of sensory transmission in the NTS. In rat second-order neurons receiving identified carotid body afferent input, α_1_ receptor activation reduced glutamatergic transmission, and subsequent experiments demonstrated contributions from both α_1_and α_2_ receptors to inhibition by NE (59, 60). The predominance of α_2_ signaling in the present study may reflect differences in afferent identity, postsynaptic phenotype, species, or experimental conditions. It is important to note that our recordings establish this mechanism only in identified mouse TH^+^ NTS neurons. The discrepancy between our results and previous works in the literature (59, 60) indicates that adrenergic modulation should be considered in the context of the particular afferent pathway and neuronal population examined.

NE increased the paired pulse ratio, which provides further functional evidence for reduced presynaptic release probability. NE preferentially inhibited the early responses within a stimulus train, consistent with a change in the initial strength and subsequent dynamics of afferent transmission. Moreover, unchanged basal sEPSC and asynchronous EPSC amplitudes also argue against a generalized reduction in postsynaptic glutamate sensitivity. The persistence of inhibition during GABA_A_ receptor blockade indicates that recruitment of GABA_A_ mediated inhibition is not required for the reduction in ST-EPSC amplitude.

α_2_ receptors act through G_i/o_ signaling, inhibiting voltage gated Ca^2+^ channels and reducing the Ca^2+^ entry available to trigger transmitter release (61, 62). Inhibition mediated by α_2_ receptors of glutamatergic transmission has also been demonstrated in other brainstem and forebrain preparations (63, 64). Such mechanisms could account for the reduced ST-EPSC amplitude and increased paired-pulse ratio observed here. However, we did not measure terminal Ca^2+^ signals or manipulate intracellular signaling pathways, so the downstream effectors responsible for α_2_-mediated NE inhibition at these synapses remain to be established.

NE also reduced the frequency of asynchronous EPSCs following ST stimulation without changing their mean amplitude. This extends its influence beyond the rapid synchronous response to the delayed glutamatergic input that follows an afferent burst. Asynchronous release is characteristic of ST afferents expressing TRPV_1_ (55, 56), but this association does not establish TRPV_1_ as the target of NE. Indeed, subsequent work distinguished synchronous and asynchronous release driven by voltage-gated Ca^2+^ channels from spontaneous release dependent on TRPV_1_ (65). The suppression of evoked and asynchronous transmission, together with the lack of a detectable effect on basal sEPSCs, is consistent with differential regulation of Ca2+ dynamics in these release modes (65). Nevertheless, spontaneous events can arise from multiple inputs, so an unchanged population average does not establish that spontaneous release is insensitive to NE at every terminal.

The response to CCK further illustrates that the absence of an effect on basal sEPSCs does not preclude modulation of facilitated excitatory input. NE reduced both the peak and sustained increases in sEPSC frequency produced by CCK. Previous work showed that CCK enhances glutamatergic afferent input to NTS catecholaminergic neurons (21), and that NPY can attenuate this response (39). Our findings extend this inhibitory modulation to NE, suggesting that noradrenergic signaling can oppose CCK-associated excitation within the NTS.

The interaction between NE and NPY offers an interesting mechanism for regulating afferent input. Previous experiments identified presynaptic Y_2_ receptors as mediators of inhibition produced by NPY at synapses onto NTS catecholaminergic neurons (39). Here, NE and NPY each reduced evoked EPSC amplitude, and their combined application produced greater inhibition than either transmitter alone. At lower concentrations, neither transmitter produced a detectable effect individually, whereas the combination produced greater inhibition than NPY alone. These findings demonstrate enhanced inhibition during combined exposure. However, the stronger combined effect also does not establish that α_2_ and Y_2_ receptors act through common or separate intracellular pathways. Both receptor systems can suppress presynaptic Ca^2+^ entry and transmitter release (61, 66, 67), and biochemical studies have documented interactions between NPY and α_2_ signaling in tissue containing the NTS (68, 69). These results may reflect incomplete recruitment of a shared pathway or modulation of different release processes. Thus, at the concentrations tested, the two transmitters do not fully occlude one another, but their subcellular relationship remains unresolved. Gabazine did not prevent the inhibition produced by either transmitter, supporting a mechanism that does not require GABA_A_ receptor activation. Combined α_2_ and Y_2_ receptor blockade had no detectable effect on basal ST transmission, indicating no measurable tonic influence under these slice *ex vivo* conditions; this does not exclude recruitment of either system in vivo.

### Physiological implications

Our findings identify presynaptic α₂ and Y₂ receptors as regulators of visceral sensory gain at the first central vagal synapse onto TH^+^ second-order NTS Neurons. By reducing synchronous, asynchronous, and CCK-evoked glutamate release, these receptors can control both the strength and duration of afferent excitation reaching NTS catecholaminergic neurons.

This mechanism may become particularly important during fasting or hypoglycemia. CCK strengthens vagal and central excitatory signaling after a meal and contributes to satiation and meal termination (21, 46, 47, 70–74). By suppressing CCK-induced glutamate release, NE may reduce the influence of postprandial satiation signals when restoring energy availability becomes a physiological priority.

However, the behavioral consequence of this presynaptic inhibition will depend on which catecholaminergic population receives the affected input. Chen et al. (2020) identified a clear functional dichotomy within the NTS: epinephrine neurons that co-express NPY promote feeding, whereas norepinephrine neurons suppress feeding. In parallel, TH^+^ NTS neurons that project to the ARC and rostral PVH promote food seeking, meal initiation, and glucoprivic feeding (20, 35). These findings indicate that catecholaminergic identity, peptide expression, and projection target together determine the behavioral output of NTS circuits. This distinction is relevant because α₂ and Y₂ receptors reduce excitatory afferent input to TH^+^ NTS neurons, but the behavioral consequence of this inhibition will depend on the identity of the postsynaptic neuron. The interaction between norepinephrine (NE) and neuropeptide Y (NPY) in suppressing ST presynaptic activity may provide a feedback mechanism through which NTS catecholaminergic (NTS-CA) neurons regulate their own excitatory input. In the context of feeding regulation, diminished glutamatergic excitation of anorexigenic catecholaminergic neurons could promote food intake, whereas reduced excitation of orexigenic catecholaminergic neurons could limit feeding. It is worth noting that these experiments were performed in ad libitum-fed animals. Because nutritional state can influence vagal afferent signaling and NTS neuronal activity (Dockray, G.J. and Burdyga, G., 2011), the effects of NE and NPY may differ depending on the animal’s feeding state, providing an additional level of regulation that could explain this divergence in activity.

Our recordings did not distinguish norepinephrine neurons from epinephrine/NPY neurons or identify their projection targets. Therefore, we cannot determine whether α₂ and Y₂ receptors preferentially regulate orexigenic or anorexigenic catecholaminergic circuits. Instead, our results identify a presynaptic mechanism that can adjust vagal afferent drive to TH^+^ NTS neurons. Future studies using phenotype and projection-defined recordings will be necessary to establish the behavioral direction of this modulation.

## Conclusion

In conclusion, our study identifies α₂ adrenergic and Y₂ receptors as complementary presynaptic regulators of vagal afferent transmission to NTS catecholaminergic neurons. NE suppresses evoked, asynchronous, and CCK-induced glutamate release through α₂ receptors, while NPY produces additional inhibition through Y₂ receptors and partially distinct intracellular mechanisms. Together, these pathways provide a powerful means of controlling the strength and duration of visceral sensory input at its first central synapse. By adjusting afferent gain according to physiological state, this mechanism may reshape how the NTS integrates satiation and energy deficit signals. The behavioral outcome will depend on whether the affected catecholaminergic neurons belong to orexigenic epinephrine/NPY circuits or anorexigenic norepinephrine circuits. Thus, presynaptic α₂ and Y₂ signaling represents a critical control point through which catecholaminergic and peptidergic systems can dynamically regulate visceral information processing and energy balance circuits.

## Data availability

All data is available upon reasonable request from Suzanne Appleyard.

## Supplemental material

## Acknowledgements

We would like to thank Crystal Dillon for her help with mice genotyping and Minyang Zhu for technical assistance with experiments. As he has retired from science, he is not included as an author.

## Grants

This work has been funded by NIH grant DK138673 (SMA).

## Disclosures

The authors of this paper have no conflict of interest to declare.

## Disclaimers

The authors of this paper have no disclaimers to declare.

## Author Contributions

Luciano G Fernandes, Josefina I Jara, Suzanne M. Appleyard All authors conceived and designed research. LGF and JIJ performed experiments. LGF, JIJ and SMA analyzed data and/or interpreted results of experiments. LGF prepared figures and drafted the initial manuscript. LGF, JIJ and SMA edited and revised the manuscript, LGF, JIJ and SMA approved final version of manuscript.

## Notes

### Competing Interest Statement

The authors have declared no competing interest.

## Reference List

1. Loewy AD. Central Autonomic Pathways. In: Central Regulation of Autonomic Functions, edited by Arthur D. Loewy KMS. Oxford University Press, 1990, p. 88–103.

2. Andresen MC, Kunze DL. Nucleus tractus solitarius--gateway to neural circulatory control. Annu Rev Physiol 56: 93–116, 1994. doi: 10.1146/annurev.ph.56.030194.000521.

3. Andresen MC, Yang MY. Non-NMDA receptors mediate sensory afferent synaptic transmission in medial nucleus tractus solitarius. Am J Physiol 259: H1307–1311, 1990. doi: 10.1152/ajpheart.1990.259.4.H1307.

4. Doyle MW, Andresen MC. Reliability of monosynaptic sensory transmission in brain stem neurons in vitro. J Neurophysiol 85: 2213–2223, 2001. doi: 10.1152/jn.2001.85.5.2213.

5. Reis DJ, Granata AR, Perrone MH, Talman WT. Evidence that glutamic acid is the neurotransmitter of baroreceptor afferent terminating in the nucleus tractus solitarius (NTS). J Auton Nerv Syst 3: 321–334, 1981. doi: 10.1016/0165-1838(81)90073-4.

6. Andresen MC, Doyle MW, Bailey TW, Jin YH. Differentiation of autonomic reflex control begins with cellular mechanisms at the first synapse within the nucleus tractus solitarius. Brazilian Journal of Medical and Biological Research 37: 549–58, 2004.

7. Spencer SE, Talman WT. Modulation of gastric and arterial pressure by nucleus tractus solitarius in rat. Am J Physiol 250: R996–1002, 1986. doi: 10.1152/ajpregu.1986.250.6.R996.

8. Talman WT, Perrone MH, Reis DJ. Evidence for L-glutamate as the neurotransmitter of baroreceptor afferent nerve fibers. Science 209: 813–815, 1980. doi: 10.1126/science.6105709.

9. Loewy AD. Central Autonomic Pathways. In: Central Regulation of Autonomic Functions, edited by Arthur D. Loewy KMS. Oxford University Press, 1990, p. 88–103.

10. Grill HJ, Hayes MR. The nucleus tractus solitarius: a portal for visceral afferent signal processing, energy status assessment and integration of their combined effects on food intake. Int J Obes (Lond) 33 Suppl 1: S11–15, 2009. doi: 10.1038/ijo.2009.10.

11. Ritter RC, Campos CA, Nasse J, Peters JH. Vagal Afferent Signaling and the Integration of Direct and Indirect Controls of Food Intake. In: Appetite and Food Intake: Central Control, edited by Harris RB. Boca Raton (FL): CRC Press/Taylor & Francis, 2017, p. 229–258.

12. Schwartz GJ. Integrative capacity of the caudal brainstem in the control of food intake. Philos Trans R Soc Lond B Biol Sci 361: 1275–1280, 2006. doi: 10.1098/rstb.2006.1862.

13. Grill HJ, Hayes MR. Hindbrain neurons as an essential hub in the neuroanatomically distributed control of energy balance. Cell Metab 16: 296–309, 2012. doi: 10.1016/j.cmet.2012.06.015.

14. Rinaman L. Ascending projections from the caudal visceral nucleus of the solitary tract to brain regions involved in food intake and energy expenditure. Brain Research 1350: 18–34, 2010.

15. Berthoud H-R. Multiple neural systems controlling food intake and body weight. Neurosci Biobehav Rev 26: 393–428, 2002. doi: 10.1016/s0149-7634(02)00014-3.

16. Browning KN, Carson KE. Central Neurocircuits Regulating Food Intake in Response to Gut Inputs-Preclinical Evidence. Nutrients 13, 2021. doi: 10.3390/nu13030908.

17. . Samson WK, Yosten GLC. A comprehensive review of the neuroscience of ingestion: the physiological control of eating: signals, neurons, and networks. Physiol Rev 102: 319–322, 2022. doi: 10.1152/physrev.00035.2021.

18. Schwartz GJ, Zeltser LM. Functional organization of neuronal and humoral signals regulating feeding behavior. Annu Rev Nutr 33: 1–21, 2013. doi: 10.1146/annurev-nutr-071812-161125.

19. Watts AG, Kanoski SE, Sanchez-Watts G, Langhans W. The physiological control of eating: signals, neurons, and networks. Physiol Rev 102: 689–813, 2022. doi: 10.1152/physrev.00028.2020.

20. . Aklan I, Sayar Atasoy N, Yavuz Y, Ates T, Coban I, Koksalar F, Filiz G, Topcu IC, Oncul M, Dilsiz P, Cebecioglu U, Alp MI, Yilmaz B, Davis DR, Hajdukiewicz K, Saito K, Konopka W, Cui H, Atasoy D. NTS Catecholamine Neurons Mediate Hypoglycemic Hunger via Medial Hypothalamic Feeding Pathways. Cell Metab 31: 313–326.e5, 2020. doi: 10.1016/j.cmet.2019.11.016.

21. . Appleyard SM, Marks D, Kobayashi K, Okano H, Low MJ, Andresen MC. Visceral afferents directly activate catecholamine neurons in the solitary tract nucleus. J Neurosci 27: 13292–13302, 2007. doi: 10.1523/JNEUROSCI.3502-07.2007.

22. Petrov T, Krukoff TL, Jhamandas JH. Branching projections of catecholaminergic brainstem neurons to the paraventricular hypothalamic nucleus and the central nucleus of the amygdala in the rat. Brain Res 609: 81–92, 1993. doi: 10.1016/0006-8993(93)90858-k.

23. Reyes BAS, Van Bockstaele EJ. Divergent projections of catecholaminergic neurons in the nucleus of the solitary tract to limbic forebrain and medullary autonomic brain regions. Brain Res 1117: 69–79, 2006. doi: 10.1016/j.brainres.2006.08.051.

24. Ritter S, Bugarith K, Dinh TT. Immunotoxic destruction of distinct catecholamine subgroups produces selective impairment of glucoregulatory responses and neuronal activation. J Comp Neurol 432: 197–216, 2001. doi: 10.1002/cne.1097.

25. Zhao Z, Wang L, Gao W, Hu F, Zhang J, Ren Y, Lin R, Feng Q, Cheng M, Ju D, Chi Q, Wang D, Song S, Luo M, Zhan C. A Central Catecholaminergic Circuit Controls Blood Glucose Levels during Stress. Neuron 95: 138–152.e5, 2017. doi: 10.1016/j.neuron.2017.05.031.

26. Mönnikes H, Lauer G, Arnold R. Peripheral administration of cholecystokinin activates c-fos expression in the locus coeruleus/subcoeruleus nucleus, dorsal vagal complex and paraventricular nucleus via capsaicin-sensitive vagal afferents and CCK-A receptors in the rat. Brain Res 770: 277–288, 1997. doi: 10.1016/s0006-8993(97)00865-2.

27. Rinaman L, Baker EA, Hoffman GE, Stricker EM, Verbalis JG. Medullary c-Fos activation in rats after ingestion of a satiating meal. Am J Physiol 275: R262–268, 1998. doi: 10.1152/ajpregu.1998.275.1.R262.

28. Willing AE, Berthoud HR. Gastric distension-induced c-fos expression in catecholaminergic neurons of rat dorsal vagal complex. Am J Physiol 272: R59–67, 1997. doi: 10.1152/ajpregu.1997.272.1.R59.

29. Palmiter RD. The Parabrachial Nucleus: CGRP Neurons Function as a General Alarm. Trends Neurosci 41: 280–293, 2018. doi: 10.1016/j.tins.2018.03.007.

30. Riche D, De Pommery J, Menetrey D. Neuropeptides and catecholamines in efferent projections of the nuclei of the solitary tract in the rat. J Comp Neurol 293: 399–424, 1990. doi: 10.1002/cne.902930306.

31. Fukuda A, Minami T, Nabekura J, Oomura Y. The effects of noradrenaline on neurones in the rat dorsal motor nucleus of the vagus, in vitro. J Physiol 393: 213– 231, 1987. doi: 10.1113/jphysiol.1987.sp016820.

32. Martinez-Peña y Valenzuela I, Rogers RC, Hermann GE, Travagli RA. Norepinephrine effects on identified neurons of the rat dorsal motor nucleus of the vagus. Am J Physiol Gastrointest Liver Physiol 286: G333–339, 2004. doi: 10.1152/ajpgi.00289.2003.

33. Sawchenko PE. Central connections of the sensory and motor nuclei of the vagus nerve. J Auton Nerv Syst 9: 13–26, 1983. doi: 10.1016/0165-1838(83)90129-7.

34. Travagli RA, Anselmi L. Vagal neurocircuitry and its influence on gastric motility. Nat Rev Gastroenterol Hepatol 13: 389–401, 2016. doi: 10.1038/nrgastro.2016.76.

35. Sayar-Atasoy N, Laule C, Aklan I, Kim H, Yavuz Y, Ates T, Coban I, Koksalar-Alkan F, Rysted J, Davis D, Singh U, Alp MI, Yilmaz B, Cui H, Atasoy D. Adrenergic modulation of melanocortin pathway by hunger signals. Nat Commun 14: 6602, 2023. doi: 10.1038/s41467-023-42362-8.

36. Rinaman L. Visceral sensory inputs to the endocrine hypothalamus. Front Neuroendocrinol 28: 50–60, 2007. doi: 10.1016/j.yfrne.2007.02.002.

37. Andresen MC, Fawley JA, Hofmann ME. Peptide and lipid modulation of glutamatergic afferent synaptic transmission in the solitary tract nucleus. Front Neurosci 6: 191, 2012. doi: 10.3389/fnins.2012.00191.

38. Appleyard SM, Bailey TW, Doyle MW, Jin Y-H, Smart JL, Low MJ, Andresen MC. Proopiomelanocortin neurons in nucleus tractus solitarius are activated by visceral afferents: regulation by cholecystokinin and opioids. J Neurosci 25: 3578– 3585, 2005. doi: 10.1523/JNEUROSCI.4177-04.2005.

39. Calkins RJ, Zhao H, Page SJ, Neyens DM, Appleyard SM. NPY inhibits vagal activation of NTS catecholamine neurons via presynaptic Y2 receptors. .

40. Cui RJ, Li X, Appleyard SM. Ghrelin inhibits visceral afferent activation of catecholamine neurons in the solitary tract nucleus. J Neurosci 31: 3484–3492, 2011. doi: 10.1523/JNEUROSCI.3187-10.2011.

41. Cui RJ, Roberts BL, Zhao H, Zhu M, Appleyard SM. Serotonin activates catecholamine neurons in the solitary tract nucleus by increasing spontaneous glutamate inputs. J Neurosci 32: 16530–16538, 2012. doi: 10.1523/JNEUROSCI.1372-12.2012.

42. Cui RJ, Roberts BL, Zhao H, Andresen MC, Appleyard SM. Opioids inhibit visceral afferent activation of catecholamine neurons in the solitary tract nucleus. Neuroscience 222: 181–190, 2012. doi: 10.1016/j.neuroscience.2012.07.010.

43. Fernandes LG, Jin YH, Andresen MC. Heterosynaptic crosstalk: GABA-glutamate metabotropic receptors interactively control glutamate release in solitary tract nucleus. Neuroscience 174: 1–9, 2011.

44. Jin Y-H, Bailey TW, Li B-Y, Schild JH, Andresen MC. Purinergic and vanilloid receptor activation releases glutamate from separate cranial afferent terminals in nucleus tractus solitarius. J Neurosci 24: 4709–4717, 2004. doi: 10.1523/JNEUROSCI.0753-04.2004.

45. Jin Y-H, Bailey TW, Andresen MC. Cranial afferent glutamate heterosynaptically modulates GABA release onto second-order neurons via distinctly segregated metabotropic glutamate receptors. J Neurosci 24: 9332–9340, 2004. doi: 10.1523/JNEUROSCI.1991-04.2004.

46. Moran TH, Ladenheim EE, Schwartz GJ. Within-meal gut feedback signaling. Int J Obes Relat Metab Disord 25 Suppl 5: S39–41, 2001. doi: 10.1038/sj.ijo.0801910.

47. Winzenried ET, Neyens DM, Calkins R, Appleyard SM. CCK-expressing neurons in the NTS are directly activated by CCK-sensitive C-type vagal afferents. Am J Physiol Regul Integr Comp Physiol 328: R121–R132, 2025. doi: 10.1152/ajpregu.00280.2023.

48. Matsushita N, Okada H, Yasoshima Y, Takahashi K, Kiuchi K, Kobayashi K. Dynamics of tyrosine hydroxylase promoter activity during midbrain dopaminergic neuron development. J Neurochem 82: 295–304, 2002. doi: 10.1046/j.1471-4159.2002.00972.x.

49. Van Rossum G, Drake FL. Python 3 Reference Manual. Scotts Valley, CA: CreateSpace, 2009.

50. Hunter JD. Matplotlib: A 2D graphics environment. Computing in Science & Engineering 9: 90–95, 2007. doi: 10.1109/MCSE.2007.55.

51. Virtanen P, Gommers R, Oliphant TE, Haberland M, Reddy T, Cournapeau D, Burovski E, Peterson P, Weckesser W, Bright J, van der Walt SJ, Brett M, Wilson J, Millman KJ, Mayorov N, Nelson ARJ, Jones E, Kern R, Larson E, Carey CJ, Polat İ, Feng Y, Moore EW, VanderPlas J, Laxalde D, Perktold J, Cimrman R, Henriksen I, Quintero EA, Harris CR, Archibald AM, Ribeiro AH, Pedregosa F, van Mulbregt P. SciPy 1.0: fundamental algorithms for scientific computing in Python. Nat Methods 17: 261–272, 2020. doi: 10.1038/s41592-019-0686-2.

52. Fawley JA, Peters JH, Andresen MC. GABA(B)-mediated inhibition of multiple modes of glutamate release in the nucleus of the solitary tract. J Neurophysiol 106: 1833–1840, 2011. doi: 10.1152/jn.00476.2011.

53. Fenwick AJ, Wu S-W, Peters JH. Isolation of TRPV1 independent mechanisms of spontaneous and asynchronous glutamate release at primary afferent to NTS synapses. Front Neurosci 8: 6, 2014. doi: 10.3389/fnins.2014.00006.

54. . Neyens DM, Brenner L, Calkins RJ, Winzenried ET, Ritter RC, Appleyard SM. CCK-sensitive C fibers activate NTS leptin receptor-expressing neurons via NMDA receptors. Am J Physiol Regul Integr Comp Physiol 326: R383–R400, 2024. doi: 10.1152/ajpregu.00238.2022.

55. Peters JH, McDougall SJ, Fawley JA, Andresen MC. TRPV1 marks synaptic segregation of multiple convergent afferents at the rat medial solitary tract nucleus. PLoS One 6: e25015, 2011. doi: 10.1371/journal.pone.0025015.

56. Peters JH, McDougall SJ, Fawley JA, Smith SM, Andresen MC. Primary afferent activation of thermosensitive TRPV1 triggers asynchronous glutamate release at central neurons. Neuron 65: 657–69, 2010.

57. Ragozzino FJ, Arnold RA, Fenwick AJ, Riley TP, Lindberg JEM, Peterson B, Peters JH. TRPM3 expression and control of glutamate release from primary vagal afferent neurons. J Neurophysiol 125: 199–210, 2021. doi: 10.1152/jn.00229.2020.

58. Wu S, Fenwick AJ, Peters JH. Channeling satiation: a primer on the role of TRP channels in the control of glutamate release from vagal afferent neurons. Physiol Behav 136: 179–184, 2014. doi: 10.1016/j.physbeh.2014.09.003.

59. Zhang W, Mifflin SW. Modulation of synaptic transmission to second-order peripheral chemoreceptor neurons in caudal nucleus tractus solitarius by alpha1-adrenoreceptors. J Pharmacol Exp Ther 320: 670–677, 2007. doi: 10.1124/jpet.106.114033.

60. Zhang W, Carreño FR, Cunningham JT, Mifflin SW. Chronic sustained hypoxia enhances both evoked EPSCs and norepinephrine inhibition of glutamatergic afferent inputs in the nucleus of the solitary tract. J Neurosci 29: 3093–3102, 2009. doi: 10.1523/JNEUROSCI.2648-08.2009.

61. Boehm S, Huck S. Inhibition of N-type calcium channels: the only mechanism by which presynaptic alpha 2-autoreceptors control sympathetic transmitter release. Eur J Neurosci 8: 1924–1931, 1996. doi: 10.1111/j.1460-9568.1996.tb01336.x.

62. Chiu K-M, Lin T-Y, Lu C-W, Wang S-J. Inhibitory effect of glutamate release from rat cerebrocortical nerve terminals by α2 adrenoceptor agonist dexmedetomidine. Eur J Pharmacol 670: 137–147, 2011. doi: 10.1016/j.ejphar.2011.08.012.

63. Bertolino M, Vicini S, Gillis R, Travagli A. Presynaptic alpha2-adrenoceptors inhibit excitatory synaptic transmission in rat brain stem. Am J Physiol 272: G654–661, 1997. doi: 10.1152/ajpgi.1997.272.3.G654.

64. Boehm S. Presynaptic alpha2-adrenoceptors control excitatory, but not inhibitory, transmission at rat hippocampal synapses. J Physiol 519 Pt 2: 439–449, 1999. doi: 10.1111/j.1469-7793.1999.0439m.x.

65. Fawley JA, Hofmann ME, Andresen MC. Distinct Calcium Sources Support Multiple Modes of Synaptic Release from Cranial Sensory Afferents. Journal of Neuroscience 36: 8957–66, 2016.

66. Qian J, Colmers WF, Saggau P. Inhibition of synaptic transmission by neuropeptide Y in rat hippocampal area CA1: modulation of presynaptic Ca2+ entry. J Neurosci 17: 8169–8177, 1997. doi: 10.1523/JNEUROSCI.17-21-08169.1997.

67. Toth PT, Bindokas VP, Bleakman D, Colmers WF, Miller RJ. Mechanism of presynaptic inhibition by neuropeptide Y at sympathetic nerve terminals. Nature 364: 635–639, 1993. doi: 10.1038/364635a0.

68. Härfstrand A, Fredholm B, Fuxe K. Inhibitory effects of neuropeptide Y on cyclic AMP accumulation in slices of the nucleus tractus solitarius region of the rat. Neurosci Lett 76: 185–190, 1987. doi: 10.1016/0304-3940(87)90713-0.

69. Härfstrand A, Fuxe K, Agnati L, Fredholm B. Reciprocal interactions between alpha 2-adrenoceptor agonist and neuropeptide Y binding sites in the nucleus tractus solitarius of the rat. A biochemic and autoradiographic analysis. J Neural Transm 75: 83–99, 1989. doi: 10.1007/BF01677422.

70. Liddle RA. Cholecystokinin cells. Annu Rev Physiol 59: 221–242, 1997. doi: 10.1146/annurev.physiol.59.1.221.

71. Liddle RA, Green GM, Conrad CK, Williams JA. Proteins but not amino acids, carbohydrates, or fats stimulate cholecystokinin secretion in the rat. Am J Physiol 251: G243–248, 1986. doi: 10.1152/ajpgi.1986.251.2.G243.

72. Moran TH, Norgren R, Crosby RJ, McHugh PR. Central and peripheral vagal transport of cholecystokinin binding sites occurs in afferent fibers. Brain Res 526: 95–102, 1990. doi: 10.1016/0006-8993(90)90253-8.

73. Roman CW, Sloat SR, Palmiter RD. A tale of two circuits: CCK(NTS) neuron stimulation controls appetite and induces opposing motivational states by projections to distinct brain regions. Neuroscience 358: 316–324, 2017. doi: 10.1016/j.neuroscience.2017.06.049.

74. Roman CW, Derkach VA, Palmiter RD. Genetically and functionally defined NTS to PBN brain circuits mediating anorexia. Nat Commun 7: 11905, 2016. doi: 10.1038/ncomms11905.

75. Chen J, Cheng M, Wang L, Zhang L, Xu D, Cao P, Wang F, Herzog H, Song S, Zhan C. A Vagal-NTS Neural Pathway that Stimulates Feeding. Curr Biol 30: 3986–3998.e5, 2020. doi: 10.1016/j.cub.2020.07.084.

